# Supplementation with Fish Oil Rich in Omega-3 Fatty Acids Delays Age-Associated Muscle Changes

**DOI:** 10.64898/2026.08.02.742308

**Authors:** Thomas Horlem, Beatriz Borges Matthesa, Diego Francis Saraiva Rodrigues, Mônica Maciel, Matheus Felipe Zazula, Luiz Claudio Fernandes, Katya Naliwaiko

**Affiliations:** Laboratório de Plasticidade Morfofuncional, Departamento de Biologia Celular, Setor de Ciências Biológicas, Universidade Federal do Paraná. Curitiba – Paraná – Brazil; Laboratório de Metabolismo Celular, Departamento de Fisiologia, Setor de Ciências Biológicas, Universidade Federal do Paraná. Curitiba – Paraná – Brazil

**Keywords:** Skeletal muscle aging, Omega-3, Early aging, Residual effects, Glucose metabolism

## Abstract

Aging of skeletal muscle is traditionally defined by progressive loss of mass and strength; however, the early events that precede these outcomes remain poorly characterized. Here, longitudinal analyses revealed that impaired glucose tolerance arises at 12 months of age in Wistar rats, before detectable changes in body composition, circulating damage markers, or muscle mass. Structural loss was preceded by functional decline and structural disorganization between 15 and 18 months. Animals exhibited marked reductions in strength, mobility, and motor coordination, accompanied by extensive remodeling of muscle architecture, including a shift toward glycolytic fiber composition, extracellular matrix expansion, reduced capillarization, and increased structural heterogeneity. Early supplementation with n-3 polyunsaturated fatty acids, initiated at midlife, significantly improved glucose tolerance, reduced adiposity, and enhanced neuromuscular performance without increasing muscle mass. These functional benefits were paralleled by reduced markers of muscle damage and attenuation of histopathological alterations, indicating preservation of tissue organization rather than hypertrophic effects. Notably, a substantial fraction of these benefits persisted after cessation of supplementation, with animals displaying sustained metabolic and structural advantages at 18 months compared to age-matched controls. Collectively, these findings support a model in which skeletal muscle aging is driven by early loss of functional and structural efficiency rather than mass decline, and demonstrate that transient nutritional intervention can durably reprogram the trajectory of muscle aging. These results highlight a critical window of intervention and position n-3 supplementation as a strategy to induce persistent resilience against age-related functional deterioration.

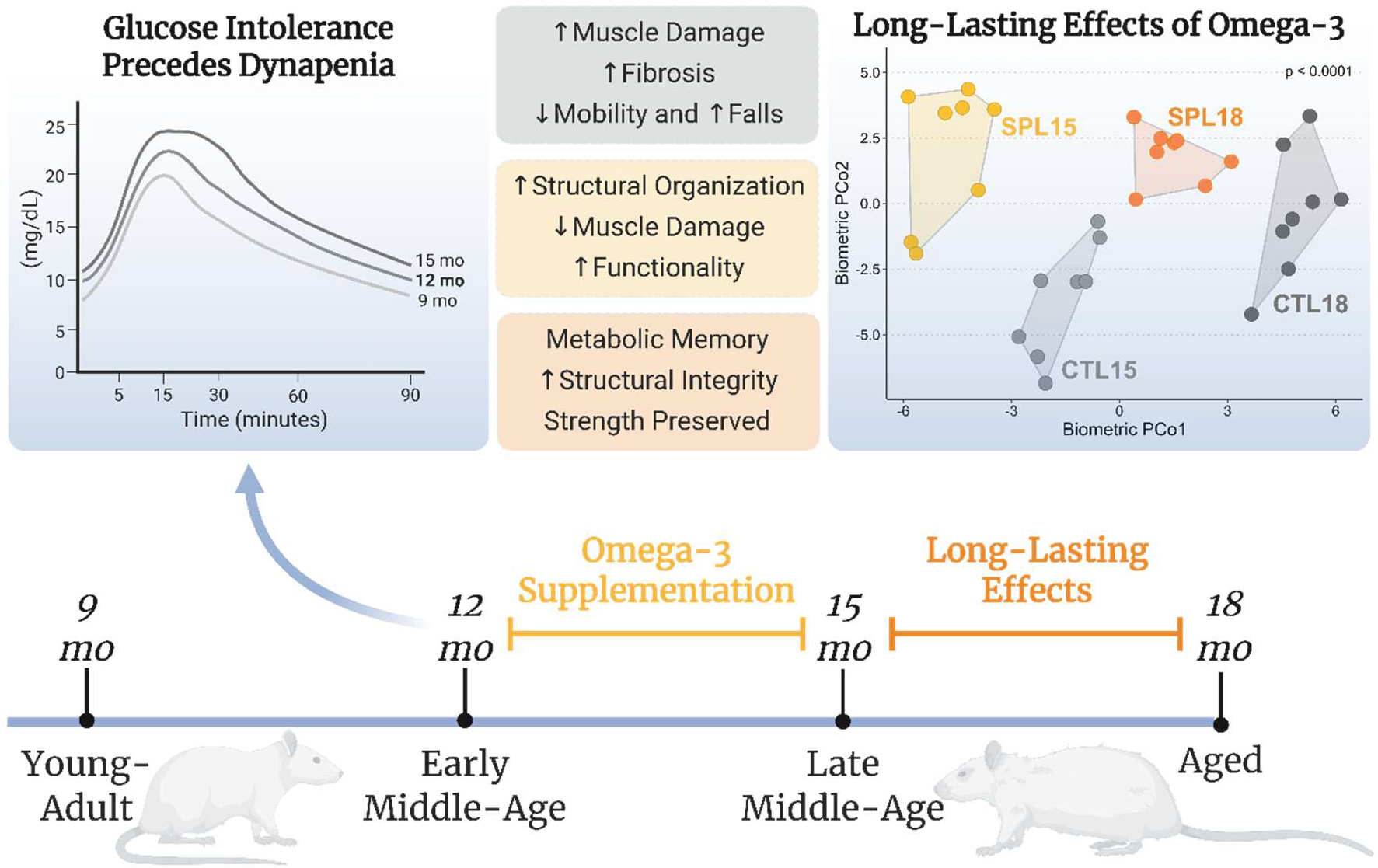

## INTRODUCTION

Population aging is advancing rapidly and transforming social, economic, and public health structures on a global scale. Estimates indicate that the number of people over 65 years of age is projected to grow to 1.6 billion by 2050, consolidating a demographic scenario in which longevity becomes a central axis of health policies [1, 2]. Investment in health during this stage of life should not be viewed merely as a cost, once it can generate broad economic benefits, including reduced healthcare expenditures and lower social security costs [3]. Therefore, strategies capable of delaying or mitigating aging-associated dysfunctions increase individual autonomy and promote social and economic benefits [3, 4].

Physiological aging involves progressive alterations associated with cellular senescence, characterized by chronic oxidative stress, loss of proteostasis, metabolic dysfunction, and reduced regenerative capacity [5]. These alterations heterogeneously affect multiple tissues, favoring the onset of aging-related chronic non-communicable diseases, such as type 2 diabetes, hypertension, cardiovascular diseases, cancer, and neurodegenerative diseases [7]. Energy metabolism is particularly vulnerable to aging. The decrease in insulin sensitivity, the increase in dysfunctional lipolysis, and the accumulation of adipose tissue contribute to glycemic worsening and damage to pancreatic β-cells [9–11].

The main tissue responsible for up to 90% of insulin-dependent glucose uptake is skeletal muscle [11]. In this sense, metabolic impacts associated with the loss of insulin sensitivity are related to various disturbances in muscle fibers, such as redox imbalance, alterations in calcium homeostasis, and a reduction in the myonuclear response, as well as in the extracellular matrix, whose deformation and stiffness with advancing age also contribute to a loss of contractile proteins and lower regenerative capacity [12–14]. This set of alterations can culminate in sarcopenia, characterized by the progressive and exponential loss of muscle mass and strength, which can reach 3% per year after age 60 [15, 16]. Negative remodeling includes fat infiltration, changes in the fiber profile, and worsening of metabolic efficiency [6, 18]. The subsequent functional losses, such as reduced mobility, balance, coordination, and gait speed, increase the risk of falls, frailty, and mortality [18, 22].

In this context, nutritional strategies have received increasing attention [15, 18]. Among them, omega-3 (n-3) polyunsaturated fatty acids, especially those derived from fish oil, demonstrate beneficial effects on energy metabolism and mitochondrial function, in addition to improving insulin sensitivity and reducing lipid accumulation in the liver, muscle, and adipose tissue [21–23]. Thus, n-3 fatty acids have also gained prominence as an adjuvant strategy in the management of muscle aging and sarcopenia [24–26]. Studies report that n-3 improves muscle protein synthesis, increases the cross-sectional area of muscle fibers, promotes contractile quality and greater strength, and enhances global functionality in older adults [26–28].

Despite this body of evidence, studies investigating the protective effect of omega-3 fatty acids on skeletal muscle prior to the onset of sarcopenia remain scarce, especially in the early stages of biological aging [31, 33]. Even more limited is the knowledge regarding the persistence of these effects after the cessation of supplementation, particularly over prolonged timeframes. From this perspective, growing evidence indicates that the first metabolic and energetic alterations in skeletal muscle appear long before the manifestation of strength loss or atrophy, characterizing a critical and underexplored window of vulnerability that begins at around 15 months of age in rats [12, 13, 32].

Given this, the present study evaluated the effects of supplementation with fish oil, rich in n-3 fatty acids (Eicosapentaenoic and Docosahexaenoic Acids - EPA and DHA), on the morphology of the plantaris muscle and the neuromuscular functional capacity of aging Wistar rats (15 months old), as well as the possible residual protective effect present months after the interruption of supplementation (18 months old).

## METHODS

### Animal Care and Experimental Groups

The Ethics Committee for Animal Use within the Biological Sciences Section of the Universidade Federal do Paraná (CEUA/BIO – UFPR) approved all animal procedures (protocol 1497/2022). These procedures were conducted in compliance with the Brazilian Guidelines for Care and Use of Animals for Scientific and Educational purposes, as established by the Conselho Nacional de Controle de Experimentação Animal (CONCEA), and adhered to international guidelines for animal research (38). Forty-eight male Wistar rats were used in this study, followed from 3 to 18 months of age. The animals were maintained with free access to standard chow (Nuvitall AR-1) and water, under a controlled temperature (23 ± 2 °C) and a 12-h light/dark cycle.

The animals were randomly allocated into two experimental groups (n = 14/group): control (CTL), with no intervention, and supplemented (SPL), which received fish oil rich in omega-3 fatty acids at a dose of 1 g·kg⁻¹·day⁻¹ orally for eight consecutive weeks between 12 and 15 months of age. Thus, the supplementation occurred in the months immediately preceding the 15-month evaluation, allowing for the assessment of immediate effects (15 months) and residual effects (18 months, three months after the end of the supplementation). The fish oil (Vitamins Sundown Naturals©) used contained 180 mg of EPA and 120 mg of DHA.

Functional and physiological evaluations were performed at the ages of 12, 15, and 18 months as described below. Blood collection for serum analysis was performed during euthanasia at 15 and 18 months; histological and histoenzymological analyses of the plantaris muscle were performed at 15 and 18 months.

### Glucose Tolerance Test

The animals were subjected to an 8-hour fast with free access to water. Glucose was administered intraperitoneally at a dose of 2 g/kg of body weight, dissolved in sterile saline solution (0.9%). The concentration of the solution was adjusted to allow a maximum injection volume of ≤ 10 mL/kg, avoiding abdominal discomfort and autonomic changes that could interfere with the glycemic response. Capillary blood glucose was measured by puncturing the distal end of the tail using a calibrated glucometer (Accu-Chek® Active, Roche). The collection times were: 0 (baseline), 15, 30, 60, and 120 min after injection. The tests were performed at 9, 12, 15, and 18 months.

### Neuromuscular Functional Evaluations

Muscle strength was evaluated using the Vertical Climbing Test, in which the animals climbed a 90 cm high vertical ladder. The protocol was employed to evaluate the functional capacity of the animal to support and move its own body weight against gravity in a vertical plane, serving as an indirect parameter of muscle performance. The execution of the test was recorded by video, and kinematic analyses were performed using the Tracker software. The videos were calibrated regarding distance in pixels, allowing the analysis of the animal’s displacement along the course. From this information, parameters of velocity, acceleration, and force developed during the test were obtained [34].

The mobility of the animals was evaluated using the Open Field Test. For this protocol, the animals were individually placed in an open-top, black arena measuring 100 cm in length, subdivided into quadrants demarcated with red adhesive tape. Each animal was filmed for a continuous period of five minutes. The positioning and camera distance parameters used for the recording were standardized according to those employed in the movement analyses. The behavioral and locomotor analysis of the videos was performed using the EthoWatcher software, responsible for quantifying the animal’s displacement in pixels throughout the test. Subsequently, a calibration between the distance covered in pixels and the actual distance in centimeters was performed, allowing the determination of the total distance covered during the experimental period. From these parameters, it was possible to evaluate and compare locomotor mobility among the different experimental groups [34].

In order to evaluate the effects of n-3 fatty acid supplementation on locomotor activity during the aging process, the Elevated Beam Walking Test was performed [34]. The protocol consisted of individually filming the animals while moving along a wooden beam 90 cm long, 1.7 cm wide, and 2 cm high, positioned between two support platforms 11 cm long and 50 cm high. After video recording the task execution, kinematic analyses were performed using the Kinovea software. The time required to cross the beam, the number of steps taken by each limb, and the number of slips that occurred during the walk were determined. From these parameters, the percentage of slips in relation to the total number of steps was calculated, allowing the evaluation of locomotor performance and motor coordination of the animals among the different experimental groups.

### Euthanasia and Tissue Collection

The animals were euthanized by decapitation; whole blood was collected and centrifuged to separate the plasma, which was stored in an ultrafreezer for biochemical analyses. The rats were weighed, and nasoanal length and tissues were collected for calculations of the Lee index, body volume, and body density.

### Metabolic and Tissue Damage Biomarkers

To assess the systemic metabolic state and lipid profile, total cholesterol, HDL, LDL, VLDL, and triacylglycerols (TAGs) were determined by colorimetric enzymatic methods in microplates. Protein status was evaluated by measuring albumin (bromocresol green method, 630 nm) and total proteins (biuret method, 540 nm). The liver enzymes aspartate aminotransferase (AST) and alanine aminotransferase (ALT) were quantified using an IFCC-standardized UV kinetic method (340 nm), while alkaline phosphatase (ALP) was determined by a colorimetric method based on the release of p-nitrophenol (405 nm).

Markers of muscle damage included total creatine kinase (total CK), CK-MM, and CK-MB, quantified by a UV kinetic method at 340 nm, based on the conversion of creatine phosphate. Lactate dehydrogenase (LDH) was determined by the same principle, monitoring the conversion of pyruvate to lactate.

All analyses were performed in triplicate using a Tecan Infinite® 200 PRO microplate reader, with validated commercial kits and according to the manufacturer’s instructions.

### Histological Processing

The plantaris muscle was dissected, coated with neutral talc powder after dissection, and kept at room temperature for approximately 15 minutes [35]. The samples were frozen in liquid nitrogen and stored in an ultrafreezer until processing. For histomorphological analysis, five-micrometer thick cross-sections of the proximal regions were acquired using a cryostat set at −25 °C (Leica, Wetzlar, Germany). The obtained sections were used for histomorphological evaluation using Hematoxylin and Eosin (H&E) staining, for collagen quantification using Picrosirius red [36, 37], and for metabolic determination by muscle fiber typing with the nicotinamide adenine dinucleotide tetrazolium reductase (NADH-TR) histoenzymological reaction [38].

Image acquisition for morphometric analysis was performed using a light microscope (Carl Zeiss™ Primo Star™). Image analysis was performed using Image-Pro Plus 6.0® software (Media Cybernetics, MD, Rockville, USA).

### Histomorphological Analysis

Morphological analysis by H&E was performed using the Histopathological Index for muscle tissue [39]. For histomorphometric analysis, images covering whole muscle fibers were used to determine the relative area of analysis, where fiber density, nuclei, and capillaries were quantified. From the data, the following parameters were calculated: Fiber density (DENS = number of fibers per mm^2^); Capillary-to-fiber ratio (CF ratio = number of capillaries/number of fibers); Nuclei-to-fiber ratio (NF ratio = number of nuclei/number of fibers); Percentage of central nuclei (CN%). The cross-sectional areas (CSA) of the muscle fibers, as well as the largest (LD) and smallest (SD) diameters of the muscle fibers and nuclei, were also measured. These measurements were employed to calculate: myonuclear domain (muscle fiber CSA/NF ratio) and the cross-sectional area ratio of myonuclei (SN CSA ratio = (NF nuclei x CSA)/fiber CSA). A total of 150 events were evaluated per animal for these measurements.

For the analysis of intramuscular collagen, slides stained with Picrosirius Red were photographed under a polarized light microscope (Carl Zeiss™ AxioImager™) and photomicrographed (Carl Zeiss™ AxioCam ERc 5s) with a camera linked to the ZEN 3.1 software (Carl Zeiss™). The quantification of thicker collagen (reddish, typically type I) and thinner collagen (greenish, typically type III) used an R script [appendix], where the pixels of the photomicrograph were quantified according to color.

### Histoenzymological Analysis

To quantify the number of type I, IIA, and IIB muscle fibers, slides treated with the NADH-TR reaction were used. All fibers in the image were counted, totaling approximately 150 fibers per animal. Subsequently, the number of each fiber type was computed, and from these data, the percentage for the proportion of each fiber type was calculated. To measure the cross-sectional area (CSA), as well as the largest (LD) and smallest (SD) diameters, 50 fibers of each type were measured per animal.

### Statistical Analysis

Data were expressed as mean ± standard deviation and analyzed using descriptive and inferential statistics in the R software (version 4.1.0). To choose the appropriate statistical test, data were evaluated for normality (Shapiro-Wilk test) and homoscedasticity (Bartlett’s test). When assumptions were met, the data were subjected to Welch’s t-test due to the inherent heteroscedasticity of aging, or otherwise, to the Wilcoxon test for nonparametric samples. The GTT analysis used a one-way ANOVA to compare glucose tolerance across ages of 9, 12, 15, and 18 months with Holm-Sidak post-hoc. In all cases, the adopted significance level was 5%.

To evaluate multivariate similarity and clustering among animals from the control and supplemented groups at 15 and 18 months of age, a Principal Coordinate Analysis (PCoA) was performed. The ordinations were conducted from three independent data matrices. The morphometric and functional matrix included the variables of nasoanal length (NAL), Lee index, retroperitoneal adipose tissue, and total adiposity, associated with metrics of physical capacity (mobility, strength, number of steps, and occurrence of falls). The serum data matrix was composed of the lipid profile (total cholesterol, triacylglycerols, HDL, LDL, and VLDL), hepatic and nutritional markers (alkaline phosphatase, aspartate aminotransferase, alanine aminotransferase, albumin, and total proteins), in addition to indicator enzymes of tissue and muscle damage (lactate dehydrogenase and total creatine kinase and its MB and MM fractions). The muscle morphology matrix included cross-sectional area (CSA), fiber proportion (types I, IIA, and IIB), diametral dimensions and their ratios, markers of tissue damage and disorganization, nuclear and myonuclear indices, as well as collagen quantification (types I, III, and mixed).

All matrices were standardized by z-score and used to calculate Euclidean distance matrices. Differences between groups were tested using Permutational Multivariate Analysis of Variance (PERMANOVA). All analyses presented degrees of freedom equal to 1 and 14 (df = 1,14). Visualization was performed through PCoA biplots, using the "ape" package and the "pcoa" function.

## RESULTS

### Onset of metabolic aging precedes overt muscular aging

The progression of metabolic aging began to be detected in the interval between 12 and 15 months, with a significant worsening in glucose tolerance between 9 and 12 months (p = 0.0333; Figure 1A), which became more pronounced at 15 and 18 months (p = 0.0005; p = 0.0207). These alterations between 15 and 18 months occurred without changes in body weight (p = 0.4980), total adiposity (p = 0.5760), or Lee index (p = 0.6710), indicating that the metabolic deterioration was not accompanied by weight gain. Muscle mass also remained stable, with no differences in the wet weight of the Tibialis anterior (p = 0.5270), soleus (p = 0.0730), and EDL (p = 0.5500) muscles. Likewise, there was no reduction in the mass, volume, or density of the plantaris muscle between 15 and 18 months (p = 0.8160; p = 0.7890; p = 0.7180; Figure 1D).

**Figure 1.**
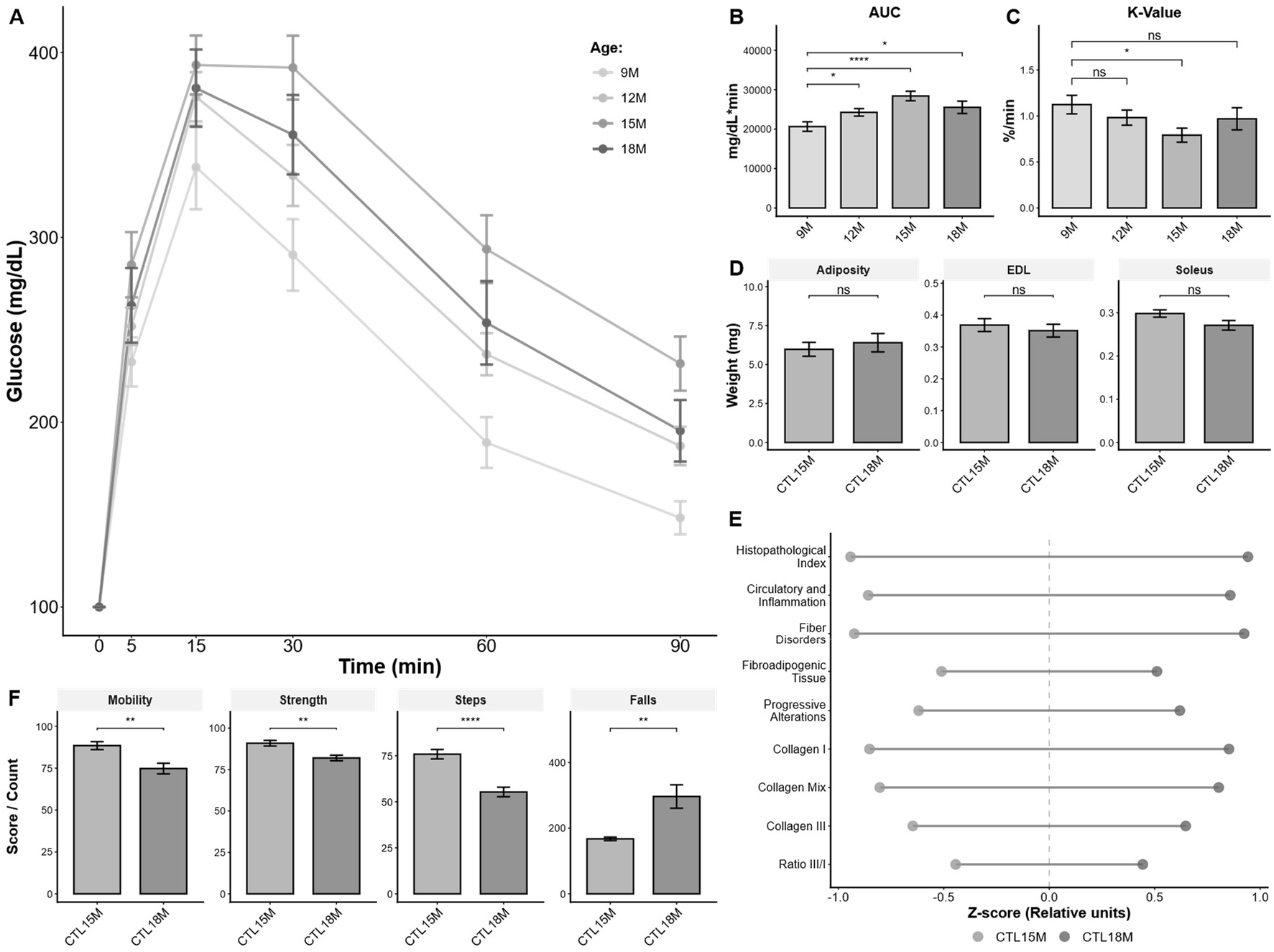
Glucose tolerance during aging in Wistar rats. (A) Temporal glycemic profile of the GTT after intraperitoneal glucose administration (2 g/kg) at different ages (9, 12, 15, and 18 months). (B) Area under the curve (AUC) of the GTT indicates glucose intolerance with age. (C) Loss of the K-value of the blood glucose clearance rate at 15 months of age. (D) Absence of changes in fat and lean body mass between CTL15M and CTL18M. (E) Disorganization of muscle architecture marks physiological aging. (F) Loss of neuromuscular functionality of the animals between 15 and 18 months.

Despite the absence of global structural alterations, there was an evident functional decline (Figure 1F). Between 15 and 18 months, the animals showed a 27% reduction in the number of steps (p ≤ 0.0001), 15% in mobility (p = 0.0044), and 10% in strength (p = 0.0023) and load carried (p = 0.0002), while falls increased by 77% (p = 0.0084), characterizing a loss of locomotor performance and motor control in early aging.

The serum profile remained largely stable between 15 and 18 months, with no alterations in the hepatic proteins ALP (p = 0.5960), AST (p = 0.1530), or ALT (p = 0.1610). Muscle damage markers also remained unchanged, including total CK (p = 0.3590), CK-MB (p = 0.9770), and CK-MM (p = 0.2390). The lipid profile showed no significant modifications for total cholesterol (p = 0.7500), TAGs (p = 0.1300), HDL (p = 0.9150), LDL (p = 0.7670), and VLDL (p = 0.1300). LDH also remained stable (p = 0.6450). The most striking serum alteration during the period was a 52% drop in albumin levels (p = 0.0001), indicating a worsening of systemic protein status.

Structural analyses of the plantaris muscle revealed consistent changes in the fiber profile. There was a reduction in the proportion of type I fibers (p = 0.0087) and an increase in type IIB fibers (p = 0.0191), while type IIA fibers remained unchanged (p = 0.4950). Type IIA fibers showed a reduction in cross-sectional area (CSA) (p = 0.0373), while type I and IIB fibers maintained their CSA (p = 0.1360; p = 0.7190). A general reduction in the largest diameter was observed (p = 0.0017), with a significant decrease in this parameter across all fiber types (I: p = 0.0001; IIA: p = 0.0002; IIB: p = 0.0027) and a tendency towards a reduction in the smallest diameter (p = 0.0664). Fiber deformation, estimated by the ratio between the largest and smallest diameters, increased mainly in type I (p = 0.0023) and IIB (p = 0.0474) fibers, with no alteration in type IIA fibers (p = 0.6340).

The muscle histopathological index demonstrated a robust aggravation of structural damage between 15 and 18 months, with an 80% increase in the total score (p ≤ 0.0001). There was an elevation in circulatory and inflammatory disturbances (p ≤ 0.0001), regressive alterations (p ≤ 0.0001), and progressive alterations (p = 0.0077), in addition to a greater accumulation of intramuscular adipose and nervous tissue (p = 0.0487). The extracellular matrix showed an increasing predominance of type I collagen (p = 0.0002), accompanied by expressive increases in type III and mixed collagen (p = 0.0002; p = 0.0009), with a tendency towards a greater proportionality of type III collagen (p = 0.0830). Capillarization per fiber decreased significantly during the period (p = 0.0401), while nuclear density increased by 27% (p = 0.0016), with a four-fold elevation in the presence of central nuclei (p ≤ 0.0001) and a 3.5-fold increase in the CSA ratio (p ≤ 0.0001).

Multivariate analysis (PCoA) evidenced a clear separation between 15- and 18-month-old animals in macroscopic (R^2^ = 0.2779; F = 5.3871; p = 0.0002) and muscular parameters (R^2^ = 0.4341; F = 10.7400; p = 0.0002), with the PCo1 axis explaining 32% and 49% of the total data variation, respectively. For serum parameters, there was no statistically significant discrimination between ages (R^2^ = 0.1130; F = 1.7834; p = 0.0838), although a greater dispersion of data was observed at 18 months, indicating an increase in physiological heterogeneity with aging (Figure 2).

**Figure 2.**
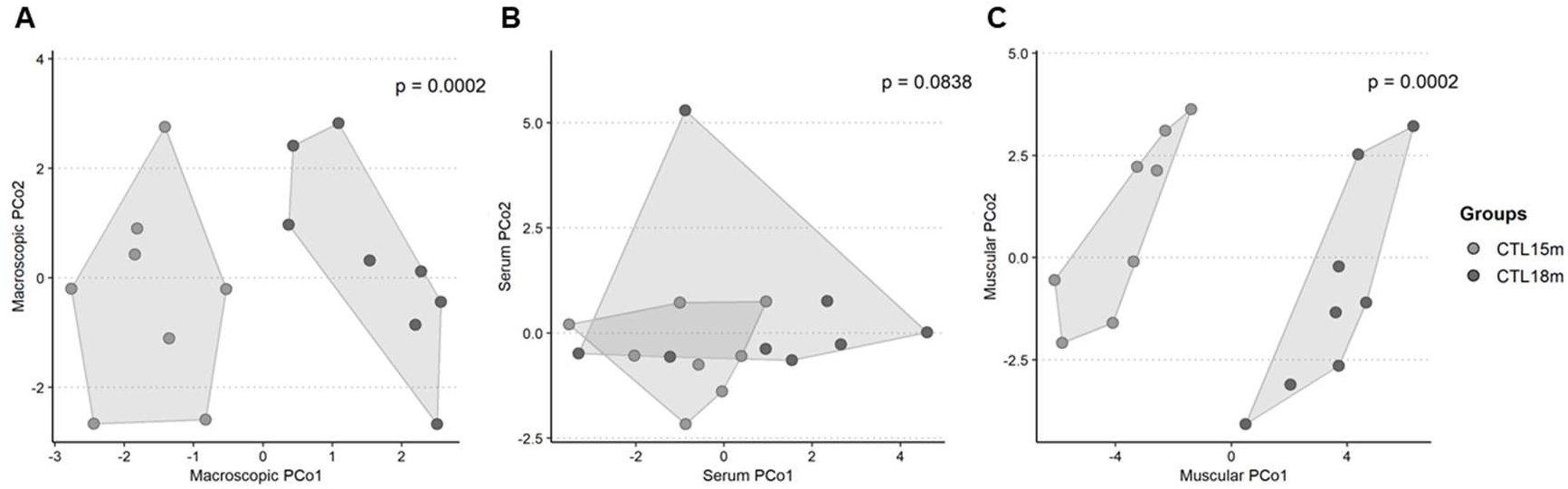
Natural aging trajectories distinctly affect the macroscopic, serum, and muscular signatures in control animals. Principal Coordinate Analysis (PCoA) plots comparing the biological profiles of control rats at 15 months (CTL15M) and 18 months (CTL18M). The panels represent the macroscopic (A), serum (B), and muscular (C) parameters. Separation between ages is observed in the macroscopic (p = 0.0002) and muscular (p = 0.0002) parameters, whereas the serum profile shows greater overlap between the groups, with no statistically significant difference (p = 0.0838). The p-values refer to the PERMANOVA analysis.

### Effect of n-3 supplementation on the aging process

Supplementation with n-3 promoted important systemic benefits at 15 months, reducing the GTT AUC (p = 0.0136; Figure 3A), although it did not alter body weight (p = 0.0521). Despite weight maintenance, a significant reduction in total adiposity was observed (p = 0.0331), resulting mainly from a decrease in mesenteric fat (p = 0.0274), with no alterations in retroperitoneal fat (p = 0.1290). The Lee index remained unchanged (p = 0.9020). Lean mass also showed no differences, including the Tibialis anterior (p = 0.9520), soleus (p = 0.4910), and EDL (p = 0.6030), and there was no effect of supplementation on the mass, volume, or density of the plantaris muscle (p = 0.8650; p = 0.1650; p = 0.5430). Despite this, musculoskeletal performance was significantly improved: there was a reduction in falls (p ≤ 0.0001) and an increase in mobility (p = 0.0002) and strength (p ≤ 0.0001), while the number of steps remained similar between groups (p = 0.0916; Figure 3F).

**Figure 3.**
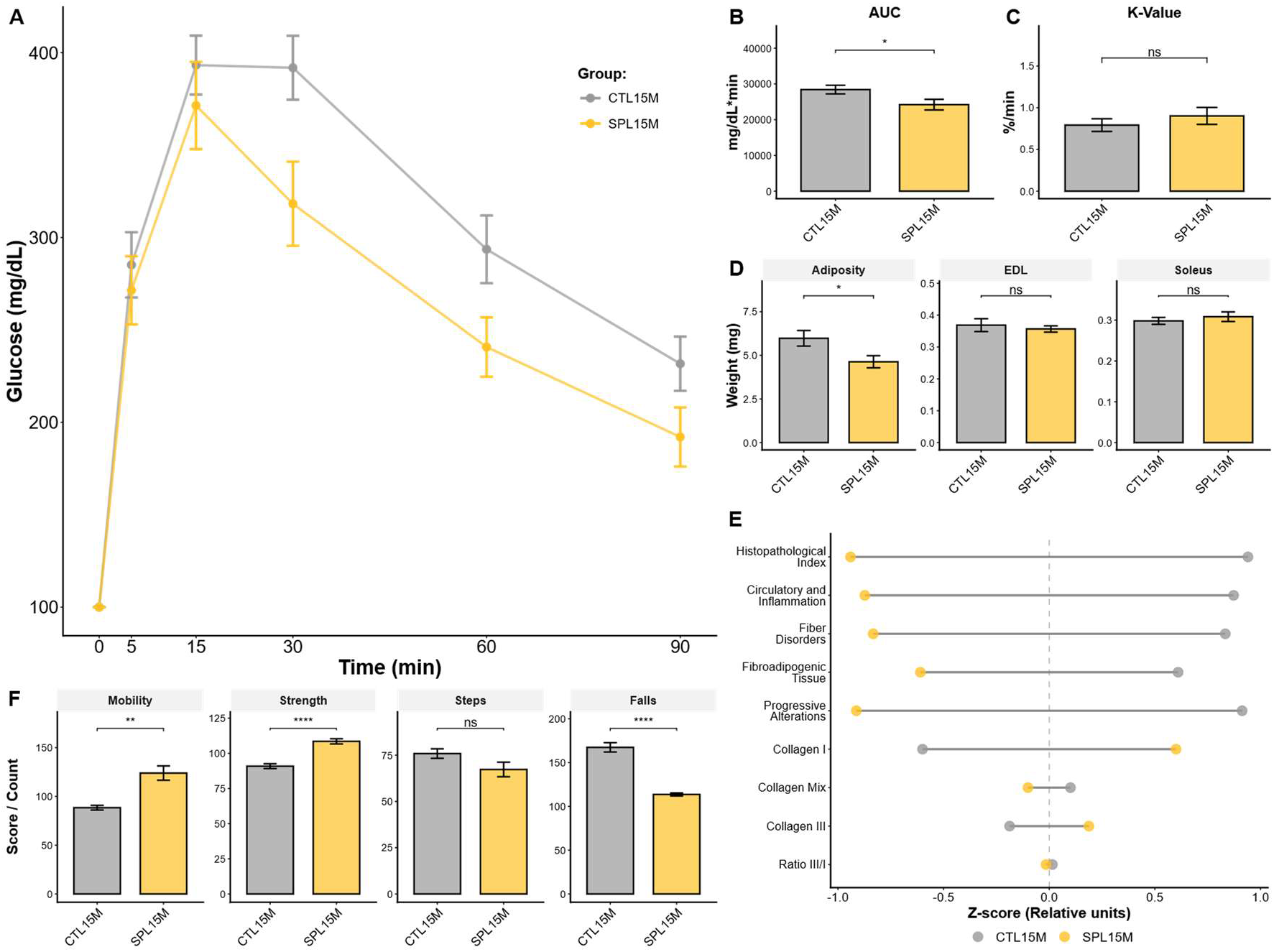
Immediate effects of omega-3 at the onset of middle age in Wistar rats. (A) Temporal glycemic profile of the GTT after intraperitoneal glucose administration (2 g/kg) in control (CTL15M) and n-3 supplemented (SPL15M) animals at 15 months of age. (B) Area under the curve (AUC) of the GTT reduced with n-3. (C) K-value of the blood glucose clearance rate after the glycemic peak. (D) Alterations in body fat mass, but not lean mass, with n-3. (E) Muscle tissue alterations delayed with n-3. (F) Functional improvement associated with the neuromuscular performance of the supplemented middle-aged animals.

In the serum profile, the hepatic markers ALP (p = 0.2790), AST (p = 0.7820), and ALT (p = 0.2340) did not differ between groups. In contrast, supplementation reduced total CK (p = 0.0065), especially due to the drop in the skeletal isoform CK-MM (p = 0.0110), while CK-MB remained stable (p = 0.5330). In lipid metabolism, TAGs (p = 0.7210) and VLDL (p = 0.7210) were not modified; however, LDL showed a significant reduction (p = 0.0030), accompanied by an increase in HDL (p = 0.0143) and a drop in total cholesterol (p = 0.0023). There was also a reduction in serum LDH (p = 0.0028). Albumin levels remained similar to those of the control group (p = 0.9990), but there was a 63% increase in total proteins (p = 0.0281).

In the plantaris muscle, supplementation increased the proportion of type IIB fibers (p = 0.0328), without altering the proportions of type I (p = 0.8030) or type IIA (p = 0.0954) fibers. There was a significant reduction in the CSA of type I (p = 0.0401), IIA (p = 0.0181), and IIB (p = 0.0130) fibers, despite the absence of alterations in the largest and smallest diameters, both overall (p = 0.8740; p = 0.3140) and specific per fiber type (I – LD: p = 0.7700, SD: p = 0.1190; IIA – LD: p = 0.5930, SD: p = 0.2280; IIB – LD: p = 0.7650, SD: p = 0.3710). Fiber shape also remained stable (I: p = 0.0914; IIA: p = 0.2620; IIB: p = 0.2880). The histopathological index demonstrated expressive improvements, including a reduction in total structural damage (p ≤ 0.0001), circulatory and inflammatory disturbances (p ≤ 0.0001), regressive alterations (p ≤ 0.0001), progressive alterations (p ≤ 0.0001), and adipose/nervous tissue accumulation (p = 0.0200). In the extracellular matrix, type I collagen showed a modest increase, approximately two-fold (p = 0.0107), while type III collagen (p = 0.7530), mixed collagen (p = 0.4620), and the proportion between collagens were not altered (p = 0.6450). Supplementation also increased the nuclei per fiber (p = 0.0136) and the frequency of central nuclei (p = 0.0099), without modifying capillarization per fiber (p = 0.8310). Thus, a 30% decrease in the myonuclear domain (p = 0.0156) and a four-fold increase in the CSA ratio (p = 0.0074) were observed.

The PCoA analysis showed a marked separation between the control and supplemented groups at 15 months (Figure 4). In the macroscopic dimension, the supplemented group exhibited an evident displacement along the PCo1 axis (45% explained variability; R^2^ = 0.3388; F = 7.1742; p = 0.0002). Similarly, the muscular PCoA revealed a clearly distinct clustering (R^2^ = 0.2802; F = 5.4496; p = 0.0005), with the PCo1 axis explaining 34% of the variation. The serum analysis, on the other hand, showed a more subtle yet significant separation (R^2^ = 0.2623; F = 4.9767; p = 0.0006), with PCoA1 and PCoA2 explaining 34% and 21% of the total variation, respectively.

**Figure 4.**
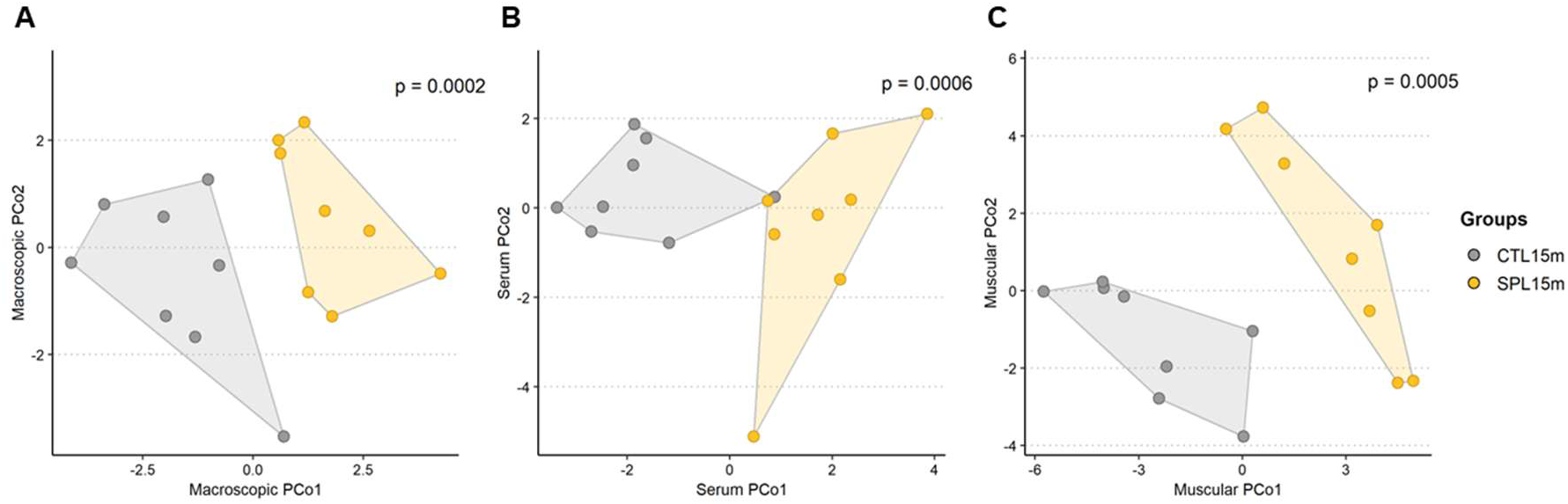
Effect of n-3 supplementation on systemic profiles at 15 months. Principal Coordinate Analysis (PCoA) plots comparing the multivariate structure between control (CTL15M) and supplemented (SPL15M) animals. Separation between the groups is observed along the principal axes for the macroscopic (A, p = 0.0002), serum (B, p = 0.0006), and muscular (C, p = 0.0005) parameters. The p-values refer to the PERMANOVA analysis.

### Residual effects of supplementation at 18 months

Even after three months without supplementation, several beneficial effects of n-3 on systemic metabolism were still present at 18 months (Figure 5). Although the mean AUC remained numerically lower than in controls, the difference was no longer statistically significant (p = 0.2786), similar identified in the adiposity data (p = 0.8030), indicating partial maintenance of metabolic protection. However, locomotor performance began to decline: falls increased significantly (p ≤ 0.0001), approaching the values observed in 15-month-old control animals. Mobility also decreased (p = 0.0002), accompanied by a reduction in the number of steps (p = 0.0098). Despite this, muscle strength remained stable between 15 and 18 months in previously supplemented animals (p = 0.6830).

**Figure 5.**
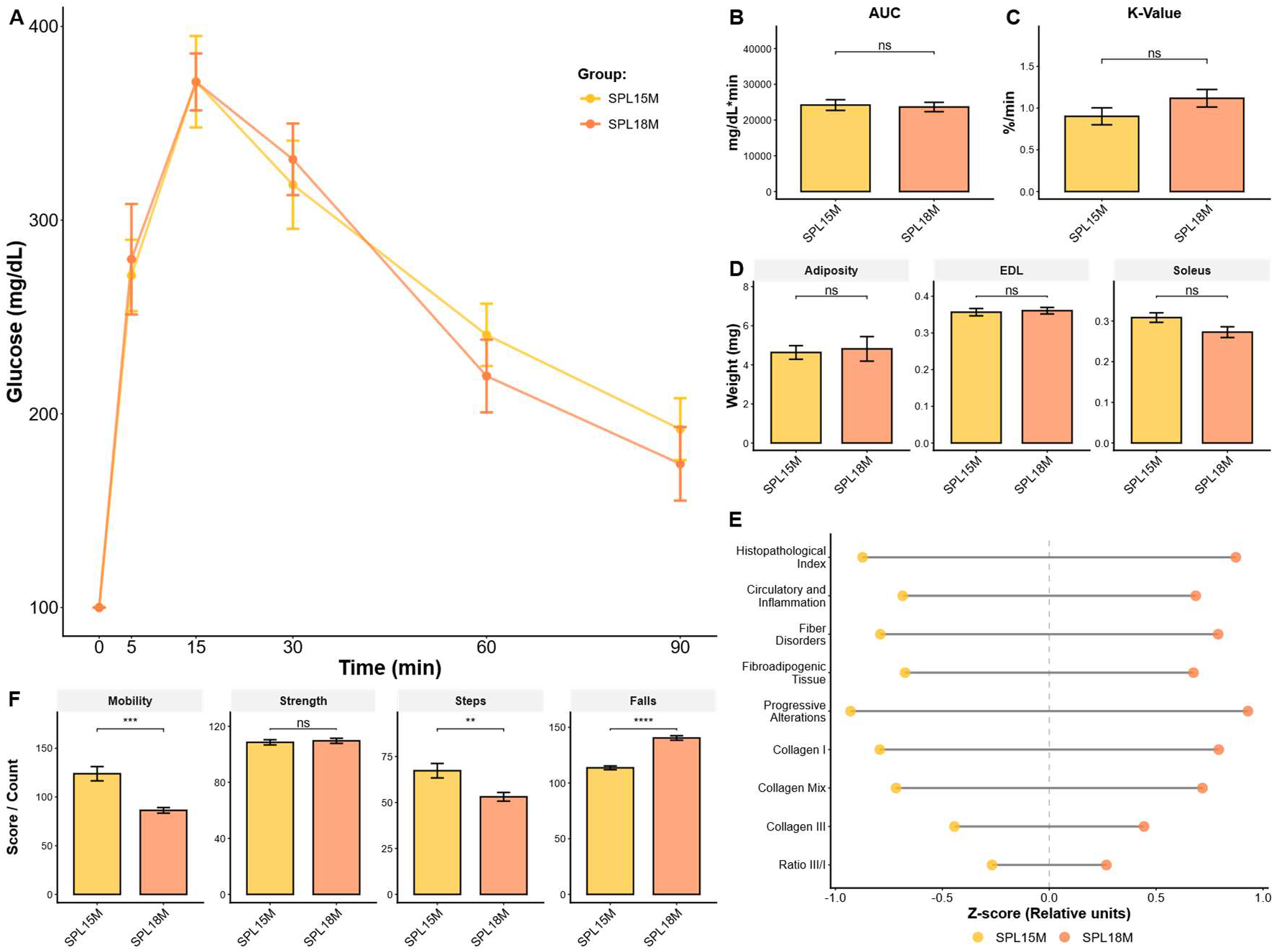
Residual effects of n-3 supplementation during aging. (A) Improvement in the temporal glycemic profile of the GTT after intraperitoneal glucose administration (2 g/kg) present at 15 months (SPL15M) remains 3 months after supplementation (SPL18M). (B) Area under the curve (AUC) of the GTT unchanged after n-3 supplementation at the onset of middle age. (C) K-value of the blood glucose clearance rate unchanged. (D) Absence of changes in fat and lean body mass between SPL15M and SPL18M. (E) Muscle tissue patterns present in aging partially resumed 3 months after supplementation. (F) Neuromuscular functional performance partially preserved between SPL15M and SPL18M.

Among the serum markers, the beneficial effects on total CK and its isoforms, CK-MM and CK-MB, were maintained even after three months without supplementation (p = 0.6430; p = 0.6030; p = 0.9750, respectively). In contrast, in the lipid profile, HDL remained unchanged (p = 0.2390), while LDL increased (p ≤ 0.0001), returning to the values observed in 15-month-old control animals. Total cholesterol remained reduced at 18 months (p = 0.3380), and the decrease in serum LDH was also maintained (p = 0.1300). However, the increase in total serum proteins observed at 15 months did not persist after the suspension of supplementation (p = 0.0281).

In the plantaris muscle structure, there was an increase in the proportion of type IIB fibers (p = 0.0404), accompanied by an increase in the CSA of these fibers (p = 0.0030). Type I and IIA fibers remained unchanged in both proportion (p = 0.0695; p = 0.6940) and CSA (p = 0.3290; p = 0.6510). Compared to animals supplemented at 15 months, an increase in the largest diameter of type I (p ≤ 0.0001), IIA (p = 0.0021), and IIB (p = 0.0100) fibers was observed, as well as an increase in the smallest diameter of IIA fibers (p = 0.0227), but not in type I (p = 0.9820) or IIB (p = 0.4510) fibers. Fiber deformation was greater in type I fibers (p = 0.0088), while type IIA (p = 0.8550) and IIB (p = 0.2010) fibers remained resistant to aging alterations even after three months without supplementation.

The positive effects of n-3 were also evident in the muscle histopathological index (Figure 5E). Although the values of total structural damage (p ≤ 0.0001), circulatory and inflammatory disturbances (p = 0.0007), regressive alterations (p = 0.0002), progressive alterations (p ≤ 0.0001), and adipose/nervous tissue accumulation (p = 0.0083) increased compared to animals supplemented at 15 months, these values were still lower than those observed in 18-month-old control animals, indicating partial protection. In the extracellular matrix, there were expressive increases after the suspension of supplementation: type I collagen rose 31-fold (p = 0.0011), type III collagen increased 98-fold (p = 0.0117), and mixed collagen increased 573-fold (p = 0.0135), although all these values remained lower than those found in 18-month-old controls. The proportion between type III and I collagens remained stable (p = 0.1720), contrary to what is observed in unmodulated aging. Nuclei per fiber (p = 0.3470) and central nuclei (p = 0.1010) remained unchanged after the period without supplementation, while the myonuclear domain (p = 0.1080) and the CSA ratio (p = 0.8780) preserved the improvements induced by n-3. However, nuclear CSA showed a significant reduction (p = 0.0057).

The PCoA analyses revealed that the residual effects of supplementation produced clear differences between supplemented and non-supplemented animals at 18 months (Figure 6). Macroscopic (R^2^ = 0.2926; F = 5.7912; p = 0.0002) and muscular (R^2^ = 0.3348; F = 7.0446; p = 0.0004) parameters sharply discriminated the groups along the PCo1 axis, which explained 40% and 37% of the total variation, respectively. Serum parameters, on the other hand, showed greater separation along the PCo2 axis (22% of the total variation), although equally significant (R^2^ = 0.2035; F = 3.5760; p = 0.0002). In general, the residual effects indicate that supplementation with n-3 PUFAs was capable of delaying the onset of alterations characteristic of physiological aging at 18 months, with the supplemented animals presenting a profile similar to that of 15-month-old controls.

**Figure 6.**
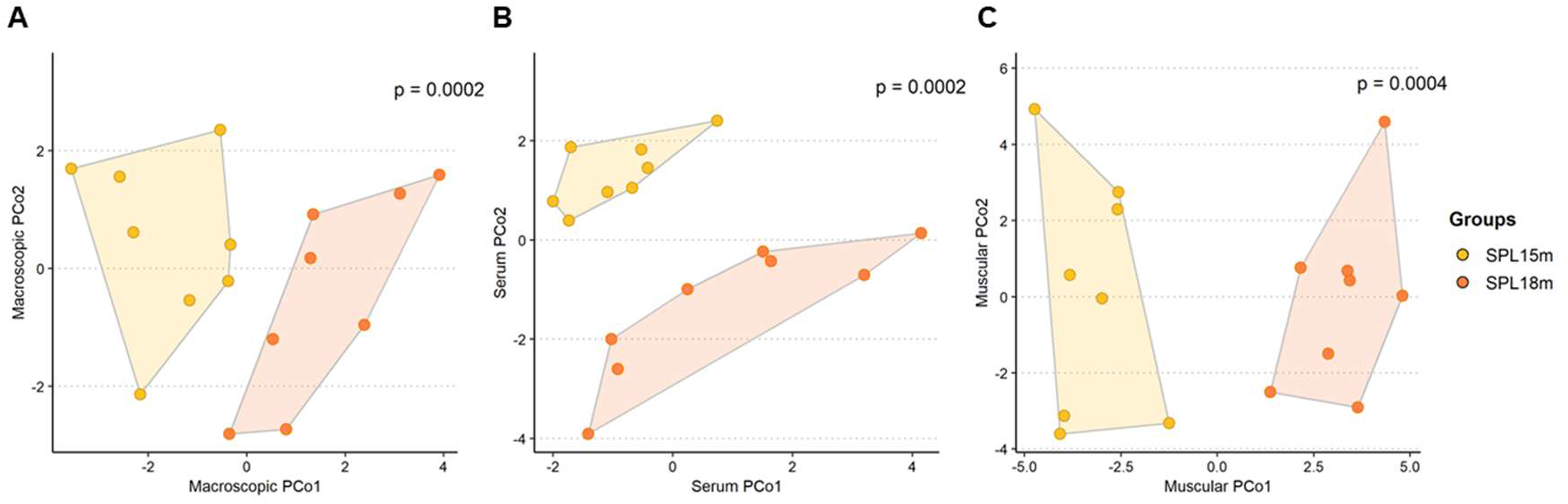
Longitudinal modulation of systemic and tissue profiles in the group even 3 months after supplementation demonstrates residual effects. Principal Coordinate Analysis (PCoA) demonstrating the distribution of phenotypic signatures in the supplemented group (SPL) at 15 and 18 months (SPL15M and SPL18M), three months after supplementation cessation. The plots show the separation between the groups along the principal axes for the macroscopic (A, p = 0.0002), serum (B, p = 0.0002), and muscular (C, p = 0.0004) parameters. The p-values indicate the statistical difference between the groups (via PERMANOVA test).

## DISCUSSION

The present study demonstrates that metabolic and muscular physiological aging in Wistar rats begins as a predominantly functional, organizational, and extracellular matrix-driven process, preceding classic quantitative alterations, such as adiposity gain, muscle mass loss, or elevation of serum damage markers. The absence of changes in body weight, total adiposity, and muscle mass up to 18 months indicates that the observed alterations do not reflect a pathological or obesogenic state, but rather a progressive loss of homeostatic efficiency characteristic of physiological aging [5, 40]. Subtle changes in the plasma profile, such as the drop in albumin concentration, reinforce that molecular disturbances precede structural alterations in the organism during aging.

In this sense, the longitudinal assessment of glucose tolerance corroborates that metabolic deterioration emerges progressively and silently [11, 41, 42]. While the tests performed at 9 months did not reveal alterations in the glycemic response, indicating the preservation of metabolic sensitivity at these stages, the evaluation at 12 months evidenced the first detectable alteration, characterized by a significant worsening of glucose tolerance. This finding establishes a timeline for the onset of metabolic aging, preceding both musculoskeletal functional decline and any measurable modification in body composition, suggesting that the loss of metabolic flexibility represents one of the first systemic signs of the physiological aging process [5, 42, 43]. Thus, our data suggest the evaluation of glucose tolerance as an early biomarker of subsequent muscle deterioration, independent of obesity, as well as serving as a parameter for the efficacy of interventions during muscle aging.

Despite the preservation of overall muscle mass, a significant functional decline was observed between 15 and 18 months, characterized by a reduction in strength, mobility, and motor coordination, as well as a significant increase in the number of falls. These findings indicate that functional impairment precedes evident structural loss, reinforcing the notion that muscle function is a more sensitive marker of aging than mass per se [49, 50]. In this context, skeletal muscle emerges not only as a target organ but as a critical integrator of the metabolic, biomechanical, and neural alterations that accompany early aging [11, 18, 52].

At the histological level, early aging was marked by a profound reorganization of muscle architecture. The reduction in the proportion of type I fibers, associated with a relative increase in type IIB fibers, indicates a transition to a more glycolytic and metabolically less efficient phenotype at the onset of aging [48]. This suggests an initial maladaptive compensatory mechanism prior to the drop in the proportion of type IIB fibers at more advanced ages [18, 19, 35].

It is important to emphasize that this redistribution occurred without a significant reduction in the overall cross-sectional area, suggesting that early aging does not involve fiber loss, but rather a change in the functional identity of the muscle reflected by early phenotypic remodeling for the preservation of glycolytic fibers [47]. Together, these data reinforce recent aging theories that point to cellular and tissue hyperfunctionality as contributors to the progressive dysfunction present in advanced age [53, 54].

In addition to the alteration in the fiber profile, changes in the geometry and shape of muscle fibers revealed a loss of structural integrity. The reduction in the largest and smallest diameters, associated with the increased deformation of the fibers, suggests an impairment in the capacity to support mechanical loads efficiently, even in the absence of evident atrophy [55, 56]. These geometric alterations, although subtle, have a direct impact on force transmission and sarcomeric stability, contributing to the observed functional decline [56]. Thus, aged muscle becomes mechanically less robust before presenting a measurable loss of mass.

Extracellular matrix remodeling stood out as a central axis of early muscle aging, reinforcing previous findings in the literature [57, 58]. The progressive increase in type I, III, and mixed collagens, accompanied by a reduction in capillarization per fiber, characterizes a more rigid, less perfused, and metabolically limiting microenvironment in aging [59–61]. This incipient fibrosis promoted by the loss of matrix collagen turnover compromises tissue elasticity, oxygen and nutrient diffusion, and paracrine communication between fibers, vessels, and satellite cells [59, 61, 62]. In this sense, the data suggest the onset of functional fibrosis that worsens at more advanced ages, in which the extracellular matrix ceases to act as adaptive support and begins to restrict muscle plasticity, preceding classic structural loss.

The increase in nuclear density and the frequency of central nuclei observed between 15 and 18 months suggests the activation of remodeling and regeneration mechanisms, possibly in response to continuous structural stress [15, 76]. However, the concomitant worsening of functional performance and the aggravation of the histopathological index indicate that these regenerative processes are insufficient or disorganized [67, 68]. Thus, the aged muscle appears to enter a state of chronic remodeling, also seen in myopathies and progerias, characterized by compensatory attempts that do not fully restore function and contribute to the structural heterogeneity of the tissue [79].

These alterations occurred in a context of relative stability of classic serum markers of muscle damage, hepatic function, and lipid metabolism, suggesting that early physiological aging does not manifest as acute cellular damage or evident systemic inflammation [6]. The significant reduction in albumin levels, however, stands out as a possible early marker of systemic vulnerability, reflecting an impairment of the protein status or long-term adaptive capacity [69, 70]. The greater dispersion observed in the multivariate analyses at 18 months reinforces this interpretation, indicating an increase in biological heterogeneity, a fundamental characteristic of aging.

Early supplementation with n-3 PUFAs significantly modulated this aging trajectory. Initiated at 12 months, the intervention promoted an improvement in glucose tolerance, a reduction in adiposity, and an expressive enhancement in musculoskeletal performance at 15 months, without inducing muscle mass gain – similar to data of n-3 effects over aged human skeletal muscle [32, 83]. These findings indicate that the beneficial effects of n-3 are predominantly associated with an increase in metabolic and functional efficiency, and not with muscle tissue expansion [72]. The reduction in serum levels of total CK, especially the CK-MM isoform, and LDH suggests less basal damage to muscle fibers and greater cell membrane stability.

In the plantaris muscle, supplementation induced a structural reorganization distinct from that observed in unmodulated aging. Although there was a reduction in the cross-sectional area of the fibers, this alteration was not accompanied by functional worsening, increased deformation, or impairment of fiber shape. These data suggest that smaller fibers, inserted in a less damaged and functionally more organized matrix microenvironment, can sustain better contractile performance [48, 73]. This dissociation between size and function reinforces the notion that the structural and metabolic quality of the muscle is more decisive for performance than its absolute volume [44, 45, 74].

The expressive improvement of the histopathological index, with a reduction in circulatory and inflammatory disturbances, regressive and progressive alterations, and the accumulation of noncontractile tissue, indicates that n-3 acts by attenuating the main vectors of structural disorganization associated with aging [75, 76]. The alterations observed in the extracellular matrix, although they include a modest increase in type I collagen, were not accompanied by a proportional increase in type III or mixed collagen, suggesting a more controlled and potentially functional remodeling of the ECM, in contrast to the progressive fibrosis observed in unmodulated aging [26, 62].

The residual effects observed after the suspension of supplementation suggest that n-3 induces phenotypic persistence throughout aging, leading to a prolonged maintenance of effects compatible with other studies that have demonstrated epigenetic and transgenerational effects of n-3 [77, 78]. Even three months after the end of the intervention, the previously supplemented animals maintained better glucose tolerance, preserved muscle strength, and less structural damage compared to controls of the same age, presenting an overall profile similar to that of younger animals. Although part of the benefits was attenuated, the deceleration of muscle aging remained evident, indicating that early supplementation alters the rate and progression pattern of aging, rather than simply delaying its effects transiently [22, 23].

Taken together, these results support a model in which physiological muscle aging begins as a process of loss of functional, metabolic, and extracellular matrix organization, prior to mass loss, and in which early nutritional interventions, such as supplementation with n-3 PUFAs, are capable of preserving systemic efficiency and delaying the consolidation of structural alterations [45, 79]. This model has relevant implications for preventive strategies in healthy aging, suggesting that the timing of the intervention is a critical determinant of its efficacy, possibly more relevant than the late attempt to reverse already established losses.

## REFERENCES

1. Navaneetham K, Arunachalam D (2025) Global Population Aging, 1950–2050. In: Handbook of Aging, Health and Public Policy. Springer, Singapore, pp 99–116

2. Xi J-Y, Lin X, Hao Y-T (2022) Measurement and projection of the burden of disease attributable to population aging in 188 countries, 1990-2050: A population-based study. J Glob Health 12:04093. 10.7189/jogh.12.04093

3. Bishop CE (2022) Economics of Aging: New Insights. J Gerontol Ser B 77:735– 738. 10.1093/geronb/gbac047

4. Beard JR, Officer A, De Carvalho IA, Sadana R, Pot AM, Michel J-P, Lloyd-Sherlock P, Epping-Jordan JE, Peeters GMEE (Geeske), Mahanani WR, Thiyagarajan JA, Chatterji S (2016) The World report on ageing and health: a policy framework for healthy ageing. The Lancet 387:2145–2154. 10.1016/S0140-6736(15)00516-4

5. López-Otín C, Blasco MA, Partridge L, Serrano M, Kroemer G (2023) Hallmarks of aging: An expanding universe. Cell 186:243–278. 10.1016/j.cell.2022.11.001

6. Ding Y, Zuo Y, Zhang B, Fan Y, Xu G, Cheng Z, Ma S, Fang S, Tian A, Gao D, Xu X, Wang Q, Jing Y, Jiang M, Xiong M, Li J, Han Z, Sun S, Wang S, He F, Yang J, Qu J, Zhang W, Liu G-H (2025) Comprehensive human proteome profiles across a 50-year lifespan reveal aging trajectories and signatures. Cell 188:5763–5784.e26. 10.1016/j.cell.2025.06.047

7. Kopp W (2024) Aging and “Age-Related” Diseases - What Is the Relation? Aging Dis 0. 10.14336/AD.2024.0570

8. Delpino FM, Figueiredo LM, Caputo EL, Mintem GC, Gigante DP (2021) What is the effect of resveratrol on obesity? A systematic review and meta-analysis. Clin Nutr ESPEN 41:59–67. 10.1016/j.clnesp.2020.11.025

9. Naito R, McKee M, Leong D, Bangdiwala S, Rangarajan S, Islam S, Yusuf S (2023) Social isolation as a risk factor for all-cause mortality: Systematic review and meta-analysis of cohort studies. PLOS ONE 18:e0280308. 10.1371/journal.pone.0280308

10. Palmer AK, Jensen MD (2022) Metabolic changes in aging humans: current evidence and therapeutic strategies. J Clin Invest 132:e158451. 10.1172/JCI158451

11. DeFronzo RA (2009) From the Triumvirate to the Ominous Octet: A New Paradigm for the Treatment of Type 2 Diabetes Mellitus. Diabetes 58:773–795. 10.2337/db09-9028

12. Lima MCDAM, Zazula MF, Martins LF, Carvalhal SR, Guimarães ATB, Fernandes LC, Naliwaiko K (2024) How soon do metabolic alterations and oxidative distress precede the reduction of muscle mass and strength in Wistar rats in aging process? Biogerontology 25:491–506. 10.1007/s10522-023-10078-3

13. Horlem T, Carvalhal SRS, Bonatto SJR, Fernandes LC (2025) Molecular Framework of the Onset and Progression of Skeletal Muscle Aging. Int J Mol Sci 26:10145. 10.3390/ijms262010145

14. Su Y, Claflin DR, Huang M, Davis CS, Macpherson PCD, Richardson A, Van Remmen H, Brooks SV (2021) Deletion of Neuronal CuZnSOD Accelerates Age-Associated Muscle Mitochondria and Calcium Handling Dysfunction That Is Independent of Denervation and Precedes Sarcopenia. Int J Mol Sci 22:10735. 10.3390/ijms221910735

15. Gras S, Blasco A, Mòdol-Caballero G, Tarabal O, Casanovas A, Piedrafita L, Barranco A, Das T, Rueda R, Pereira SL, Navarro X, Esquerda JE, Calderó J (2021) Beneficial effects of dietary supplementation with green tea catechins and cocoa flavanols on aging-related regressive changes in the mouse neuromuscular system. Aging 13:18051–18093. 10.18632/aging.203336

16. Larsson L, Degens H, Li M, Salviati L, Lee YI, Thompson W, Kirkland JL, Sandri M (2019) Sarcopenia: Aging-Related Loss of Muscle Mass and Function. Physiol Rev 99:427–511. 10.1152/physrev.00061.2017

17. Nilwik R, Snijders T, Leenders M, Groen BBL, Van Kranenburg J, Verdijk LB, Van Loon LJC (2013) The decline in skeletal muscle mass with aging is mainly attributed to a reduction in type II muscle fiber size. Exp Gerontol 48:492–498. 10.1016/j.exger.2013.02.012

18. Chen L-K (2023) Unveiling the hidden epidemic: Anorexia of aging and nutritional decline in older adults. Arch Gerontol Geriatr 111:105064. 10.1016/j.archger.2023.105064

19. Simpson RJ, Pawelec G (2021) Is mechanical loading essential for exercise to preserve the aging immune system? Immun Ageing 18:26, s12979–021-00238–9. 10.1186/s12979-021-00238-9

20. Ladang A, Beaudart C, Reginster J-Y, Al-Daghri N, Bruyère O, Burlet N, Cesari M, Cherubini A, Da Silva MC, Cooper C, Cruz-Jentoft AJ, Landi F, Laslop A, Maggi S, Mobasheri A, Ormarsdottir S, Radermecker R, Visser M, Yerro MCP, Rizzoli R, Cavalier E (2023) Biochemical Markers of Musculoskeletal Health and Aging to be Assessed in Clinical Trials of Drugs Aiming at the Treatment of Sarcopenia: Consensus Paper from an Expert Group Meeting Organized by the European Society for Clinical and Economic Aspects of Osteoporosis, Osteoarthritis and Musculoskeletal Diseases (ESCEO) and the Centre Académique de Recherche et d’Expérimentation en Santé (CARES SPRL), Under the Auspices of the World Health Organization Collaborating Center for the Epidemiology of Musculoskeletal Conditions and Aging. Calcif Tissue Int 112:197–217. 10.1007/s00223-022-01054-z

21. McGlory C, Calder PC, Nunes EA (2019) The Influence of Omega-3 Fatty Acids on Skeletal Muscle Protein Turnover in Health, Disuse, and Disease. Front Nutr 6:144. 10.3389/fnut.2019.00144

22. Xiong Y, Li X, Liu J, Luo P, Zhang H, Zhou H, Ling X, Zhang M, Liang Y, Chen Q, Xing C, Li F, Miao J, Shen W, Zhou S, Wang X, Hou FF, Liu Y, Ma K, Zhao AZ, Zhou L (2024) Omega-3 PUFAs slow organ aging through promoting energy metabolism. Pharmacol Res 208:107384. 10.1016/j.phrs.2024.107384

23. Bischoff-Ferrari HA, Gängler S, Wieczorek M, Belsky DW, Ryan J, Kressig RW, Stähelin HB, Theiler R, Dawson-Hughes B, Rizzoli R, Vellas B, Rouch L, Guyonnet S, Egli A, Orav EJ, Willett W, Horvath S (2025) Individual and additive effects of vitamin D, omega-3 and exercise on DNA methylation clocks of biological aging in older adults from the DO-HEALTH trial. Nat Aging 5:376–385. 10.1038/s43587-024-00793-y

24. Dalle C, Ostermann AI, Konrad T, Coudy-Gandilhon C, Decourt A, Barthélémy J-C, Roche F, Féasson L, Mazur A, Béchet D, Schebb NH, Gladine C (2019) Muscle Loss Associated Changes of Oxylipin Signatures During Biological Aging: An Exploratory Study From the PROOF Cohort. J Gerontol Ser A 74:608–615. 10.1093/gerona/gly187

25. Okamura T, Hashimoto Y, Miki A, Kaji A, Sakai R, Iwai K, Osaka T, Ushigome E, Hamaguchi M, Yamazaki M, Fukui M (2020) Reduced dietary omega-3 fatty acids intake is associated with sarcopenia in elderly patients with type 2 diabetes: a cross-sectional study of KAMOGAWA-DM cohort study. J Clin Biochem Nutr 66:233–237. 10.3164/jcbn.19-85

26. Del Re FM, Russ DW, Dimova KP, Scordilis SP (2025) Proteomes of aging and omega-3 supplementation in rat soleus skeletal muscle. PLOS One 20:e0323602. 10.1371/journal.pone.0323602

27. Tseng P-T, Zeng B-Y, Zeng B-S, Liao Y-C, Stubbs B, Kuo JS, Sun C-K, Cheng Y-S, Chen Y-W, Chen T-Y, Tu Y-K, Lin P-Y, Hsu C-W, Li D-J, Liang C-S, Suen M-W, Wu Y-C, Shiue Y-L, Su K-P (2023) Omega-3 polyunsaturated fatty acids in sarcopenia management: A network meta-analysis of randomized controlled trials. Ageing Res Rev 90:102014. 10.1016/j.arr.2023.102014

28. Huang Y-H, Chiu W-C, Hsu Y-P, Lo Y-L, Wang Y-H (2020) Effects of Omega-3 Fatty Acids on Muscle Mass, Muscle Strength and Muscle Performance among the Elderly: A Meta-Analysis. Nutrients 12:3739. 10.3390/nu12123739

29. Cruz-Jentoft AJ, Bahat G, Bauer J, Boirie Y, Bruyère O, Cederholm T, Cooper C, Landi F, Rolland Y, Sayer AA, Schneider SM, Sieber CC, Topinkova E, Vandewoude M, Visser M, Zamboni M, Writing Group for the European Working Group on Sarcopenia in Older People 2 (EWGSOP2), and the Extended Group for EWGSOP2, Bautmans I, Baeyens J-P, Cesari M, Cherubini A, Kanis J, Maggio M, Martin F, Michel J-P, Pitkala K, Reginster J-Y, Rizzoli R, Sánchez-Rodríguez D, Schols J (2019) Sarcopenia: revised European consensus on definition and diagnosis. Age Ageing 48:16–31. 10.1093/ageing/afy169

30. Santos DNDD, Coelho CG, Diniz MDFHS, Duncan BB, Schmidt MI, Bensenor IJM, Szlejf C, Telles RW, Barreto SM (2024) Dynapenia and sarcopenia: association with the diagnosis, duration and complication of type 2 diabetes mellitus in ELSA-Brasil. Cad Saúde Pública 40:e00081223. 10.1590/0102-311xen081223

31. Huang J, He F, Gu X, Chen S, Tong Z, Zhong S (2021) Estimation of sarcopenia prevalence in individuals at different ages from Zheijang province in China. Aging 13:6066–6075. 10.18632/aging.202567

32. Del Campo A, Contreras-Hernández I, Castro-Sepúlveda M, Campos CA, Figueroa R, Tevy MF, Eisner V, Casas M, Jaimovich E (2018) Muscle function decline and mitochondria changes in middle age precede sarcopenia in mice. Aging 10:34–55. 10.18632/aging.101358

33. Percie Du Sert N, Hurst V, Ahluwalia A, Alam S, Avey MT, Baker M, Browne WJ, Clark A, Cuthill IC, Dirnagl U, Emerson M, Garner P, Holgate ST, Howells DW, Karp NA, Lazic SE, Lidster K, MacCallum CJ, Macleod M, Pearl EJ, Petersen OH, Rawle F, Reynolds P, Rooney K, Sena ES, Silberberg SD, Steckler T, Würbel H (2020) The ARRIVE guidelines 2.0: Updated guidelines for reporting animal research. Br J Pharmacol 177:3617–3624. 10.1111/bph.15193

34. Knorr S, Rauschenberger L, Lang T, Volkmann J, Ip CW (2021) Multifactorial Assessment of Motor Behavior in Rats after Unilateral Sciatic Nerve Crush Injury. J Vis Exp 62606. 10.3791/62606

35. Cavallotti C, Mione MC, Napoleone P, Amenta F (1984) Protocol for improving the morphology of frozen sections of nervous and muscular tissue. Ital J Neurol Sci 5:99–99. 10.1007/BF02043979

36. Junqueira LCU, Bignolas G, Brentani RR (1979) Picrosirius staining plus polarization microscopy, a specific method for collagen detection in tissue sections. Histochem J 11:447–455. 10.1007/BF01002772

37. Kammerer J, Cirnu A, Williams T, Hasselmeier M, Nörpel M, Chen R, Gerull B (2024) Macro-based collagen quantification and segmentation in picrosirius red-stained heart sections using light microscopy. Biol Methods Protoc 9:bpae027. 10.1093/biomethods/bpae027

38. Tsairis P (1974) Muscle Biopsy: A Modern Approach. Arch Neurol 31:143–143. 10.1001/archneur.1974.00490380091018

39. Zazula MF, De Andrade BZ, Toni Boaro CD, Kirsch CB, Reginato A, Peretti AL, Costa RM, Bertolini GRF, Naliwaiko K, Guimarães ATB, Chasko Ribeiro LDF (2022) Development of a histopathological index for skeletal muscle analysis in Rattus norvegicus (Rodentia: Muridae). Acta Histochem 124:151892. 10.1016/j.acthis.2022.151892

40. Taffet GE (2024) Physiology of Aging. In: Wasserman MR, Bakerjian D, Linnebur S, Brangman S, Cesari M, Rosen S (eds) Geriatric Medicine. Springer International Publishing, Cham, pp 1555–1565

41. Escrivá F, Gavete ML, Fermín Y, Pérez C, Gallardo N, Alvarez C, Andrés A, Ros M, Carrascosa JM (2007) Effect of age and moderate food restriction on insulin sensitivity in Wistar rats: role of adiposity. J Endocrinol 194:131–141. 10.1677/joe.1.07043

42. Curl CC, Leija RG, Arevalo JA, Osmond AD, Duong JJ, Huie MJ, Masharani U, Horning MA, Brooks GA (2024) Altered glucose kinetics occurs with aging: a new outlook on metabolic flexibility. Am J Physiol-Endocrinol Metab 327:E217–E228. 10.1152/ajpendo.00091.2024

43. Shoemaker ME, Pereira SL, Mustad VA, Gillen ZM, McKay BD, Lopez-Pedrosa JM, Rueda R, Cramer JT (2022) Differences in muscle energy metabolism and metabolic flexibility between sarcopenic and nonsarcopenic older adults. J Cachexia Sarcopenia Muscle 13:1224–1237. 10.1002/jcsm.12932

44. Goodpaster BH, Park SW, Harris TB, Kritchevsky SB, Nevitt M, Schwartz AV, Simonsick EM, Tylavsky FA, Visser M, Newman AB, for the Health ABC Study (2006) The Loss of Skeletal Muscle Strength, Mass, and Quality in Older Adults: The Health, Aging and Body Composition Study. J Gerontol A Biol Sci Med Sci 61:1059–1064. 10.1093/gerona/61.10.1059

45. Isanejad M, Tajik B, McArdle A, Tuppurainen M, Sirola J, Kröger H, Rikkonen T, Erkkilä A (2022) Dietary omega-3 polyunsaturated fatty acid and alpha-linolenic acid are associated with physical capacity measure but not muscle mass in older women 65–72 years. Eur J Nutr 61:1813–1821. 10.1007/s00394-021-02773-z

46. Shero JA, Lindholm ME, Sandri M, Stanford KI (2025) Skeletal Muscle as a Mediator of Interorgan Crosstalk During Exercise: Implications for Aging and Obesity. Circ Res 136:1407–1432. 10.1161/CIRCRESAHA.124.325614

47. Akasaki Y, Ouchi N, Izumiya Y, Bernardo BL, LeBrasseur NK, Walsh K (2014) Glycolytic fast-twitch muscle fiber restoration counters adverse age-related changes in body composition and metabolism. Aging Cell 13:80–91. 10.1111/acel.12153

48. Van Wessel T, De Haan A, Van Der Laarse WJ, Jaspers RT (2010) The muscle fiber type–fiber size paradox: hypertrophy or oxidative metabolism? Eur J Appl Physiol 110:665–694. 10.1007/s00421-010-1545-0

49. Lee W-S, Cheung W-H, Qin L, Tang N, Leung K-S (2006) Age-associated Decrease of Type IIA/B Human Skeletal Muscle Fibers: Clin Orthop 450:231–237. 10.1097/01.blo.0000218757.97063.21

50. Zhang F-M, Wu H-F, Wang K-F, Yu D-Y, Zhang X-Z, Ren Q, Chen W-Z, Lin F, Yu Z, Zhuang C-L (2024) Transcriptome profiling of fast/glycolytic and slow/oxidative muscle fibers in aging and obesity. Cell Death Dis 15:459. 10.1038/s41419-024-06851-y

51. Horwath O, Moberg M, Edman S, Philp A, Apró W (2025) Ageing leads to selective type II myofibre deterioration and denervation independent of reinnervative capacity in human skeletal muscle. Exp Physiol 110:277–292. 10.1113/EP092222

52. Sadaki S, Tsuji R, Hayashi T, Watanabe M, Iwai R, Wenchao G, Semenova EA, Sultanov RI, Zhelankin AV, Generozov EV, Ahmetov II, Sakakibara I, Ojima K, Sakurai H, Muratani M, Kudo T, Takahashi S, Fujita R (2025) Large MAF transcription factors reawaken evolutionarily dormant fast-glycolytic type IIb myofibers in human skeletal muscle. Skelet Muscle 15:19. 10.1186/s13395-025-00391-5

53. Gems D (2022) The hyperfunction theory: An emerging paradigm for the biology of aging. Ageing Res Rev 74:101557. 10.1016/j.arr.2021.101557

54. Blagosklonny MV (2022) Cell senescence, rapamycin and hyperfunction theory of aging. Cell Cycle 21:1456–1467. 10.1080/15384101.2022.2054636

55. Lieber RL, Ward SR (2011) Skeletal muscle design to meet functional demands. Philos Trans R Soc B Biol Sci 366:1466–1476. 10.1098/rstb.2010.0316

56. Soendenbroe C, Karlsen A, Svensson RB, Kjaer M, Andersen JL, Mackey AL (2024) Marked irregular myofiber shape is a hallmark of human skeletal muscle ageing and is reversed by heavy resistance training. J Cachexia Sarcopenia Muscle 15:306–318. 10.1002/jcsm.13405

57. Lukjanenko L, Karaz S, Stuelsatz P, Gurriaran-Rodriguez U, Michaud J, Dammone G, Sizzano F, Mashinchian O, Ancel S, Migliavacca E, Liot S, Jacot G, Metairon S, Raymond F, Descombes P, Palini A, Chazaud B, Rudnicki MA, Bentzinger CF, Feige JN (2019) Aging Disrupts Muscle Stem Cell Function by Impairing Matricellular WISP1 Secretion from Fibro-Adipogenic Progenitors. Cell Stem Cell 24:433–446.e7. 10.1016/j.stem.2018.12.014

58. Duran-Jimenez B, Dobler D, Moffatt S, Rabbani N, Streuli CH, Thornalley PJ, Tomlinson DR, Gardiner NJ (2009) Advanced Glycation End Products in Extracellular Matrix Proteins Contribute to the Failure of Sensory Nerve Regeneration in Diabetes. Diabetes 58:2893–2903. 10.2337/db09-0320

59. Kragstrup TW, Kjaer M, Mackey AL (2011) Structural, biochemical, cellular, and functional changes in skeletal muscle extracellular matrix with aging. Scand J Med Sci Sports 21:749–757. 10.1111/j.1600-0838.2011.01377.x

60. Rodrigues CJ, Rodrigues, Jr. AJ, Bohm GM (1996) Effects of Aging on Muscle Fibers and Collagen Content of the Diaphragm: A Comparison with the Rectus abdominis Muscle. Gerontology 42:218–228. 10.1159/000213796

61. Taketa Y, Takahashi H (2026) Intramuscular collagen accumulation in different types of skeletal muscle fibers in middle-aged male rats. J Toxicol Pathol 39:45–50. 10.1293/tox.2025-0072

62. Kanazawa Y, Miyachi R, Higuchi T, Sato H (2023) Effects of Aging on Collagen in the Skeletal Muscle of Mice. Int J Mol Sci 24:13121. 10.3390/ijms241713121

63. Blau HM, Cosgrove BD, Ho ATV (2015) The central role of muscle stem cells in regenerative failure with aging. Nat Med 21:854–862. 10.1038/nm.3918

64. Sousa NS, Bica M, Brás MF, Sousa AC, Antunes IB, Encarnação IA, Costa TM, Martins IB, Barbosa-Morais NL, Sousa-Victor P, Neves J (2024) The immune landscape of murine skeletal muscle regeneration and aging. Cell Rep 43:114975. 10.1016/j.celrep.2024.114975

65. GrönholdtKlein M, Gorzi A, Wang L, Edström E, Rullman E, Altun M, Ulfhake B (2023) Emergence and Progression of Behavioral Motor Deficits and Skeletal Muscle Atrophy across the Adult Lifespan of the Rat. Biology 12:1177. 10.3390/biology12091177

66. Riparini G, Mackenzie M, Naz F, Brooks S, Jiang K, Deewan A, Dulek B, Islam S, Ko KD, Tsai WL, Gadina M, Dell’Orso S, Sartorelli V (2025) Epigenetic dysregulation in aged muscle stem cells drives mesenchymal progenitor expansion via IL-6 and Spp1 signaling. Nat Aging 5:2399–2416. 10.1038/s43587-025-01002-0

67. Kurland JV, Cutler AA, Stanley JT, Betta ND, Van Deusen A, Pawlikowski B, Hall M, Antwine T, Russell A, Allen MA, Dowell R, Olwin B (2023) Aging disrupts gene expression timing during muscle regeneration. Stem Cell Rep 18:1325–1339. 10.1016/j.stemcr.2023.05.005

68. Corbu A, Scaramozza A, Badiali-DeGiorgi L, Tarantino L, Papa V, Rinaldi R, D’Alessandro R, Zavatta M, Laus M, Lattanzi G, Cenacchi G (2010) Satellite cell characterization from aging human muscle. Neurol Res 32:63–72. 10.1179/174313209X385725

69. Gomi I, Fukushima H, Shiraki M, Miwa Y, Ando T, Takai K, Moriwaki H (2007) Relationship between Serum Albumin Level and Aging in Community-Dwelling Self-Supported Elderly Population. J Nutr Sci Vitaminol (Tokyo) 53:37–42. 10.3177/jnsv.53.37

70. Salive ME, Cornoni-Huntley J, Phillips CL, Guralnik JM, Cohen HJ, Ostfeld AM, Wallace RB (1992) Serum albumin in older persons: Relationship with age and health status. J Clin Epidemiol 45:213–221. 10.1016/0895-4356(92)90081-W

71. Kunz HE, Michie KL, Gries KJ, Zhang X, Ryan ZC, Lanza IR (2022) A Randomized Trial of the Effects of Dietary n3-PUFAs on Skeletal Muscle Function and Acute Exercise Response in Healthy Older Adults. Nutrients 14:3537. 10.3390/nu14173537

72. Smith GI, Julliand S, Reeds DN, Sinacore DR, Klein S, Mittendorfer B (2015) Fish oil–derived n−3 PUFA therapy increases muscle mass and function in healthy older adults. Am J Clin Nutr 102:115–122. 10.3945/ajcn.114.105833

73. Tumasian RA, Harish A, Kundu G, Yang J-H, Ubaida-Mohien C, Gonzalez-Freire M, Kaileh M, Zukley LM, Chia CW, Lyashkov A, Wood WH, Piao Y, Coletta C, Ding J, Gorospe M, Sen R, De S, Ferrucci L (2021) Skeletal muscle transcriptome in healthy aging. Nat Commun 12:2014. 10.1038/s41467-021-22168-2

74. Mitchell WK, Williams J, Atherton P, Larvin M, Lund J, Narici M (2012) Sarcopenia, Dynapenia, and the Impact of Advancing Age on Human Skeletal Muscle Size and Strength; a Quantitative Review. Front Physiol 3:. 10.3389/fphys.2012.00260

75. Fountain WA, Naruse M, Claiborne A, Trappe S, Trappe TA (2023) Controlling Inflammation Improves Aging Skeletal Muscle Health. Exerc Sport Sci Rev 51:51–56. 10.1249/JES.0000000000000313

76. Draganidis D, Jamurtas AZ, Chondrogianni N, Mastorakos G, Jung T, Grune T, Papadopoulos C, Papanikolaou K, Papassotiriou I, Papaevgeniou N, Poulios A, Batrakoulis A, Deli CK, Georgakouli K, Chatzinikolaou A, Karagounis LG, Fatouros IG (2021) Low-Grade Systemic Inflammation Interferes with Anabolic and Catabolic Characteristics of the Aged Human Skeletal Muscle. Oxid Med Cell Longev 2021:8376915. 10.1155/2021/8376915

77. Coluccia A, Borracci P, Renna G, Giustino A, Latronico T, Riccio P, Carratù MR (2009) Developmental omega-3 supplementation improves motor skills in juvenile-adult rats. Int J Dev Neurosci 27:599–605. 10.1016/j.ijdevneu.2009.05.011

78. Andrade-da-Costa BLDS, Isaac AR, Augusto RL, De Souza RF, Freitas HR, De Melo Reis RA (2019) Epigenetic Effects of Omega-3 Fatty Acids on Neurons and Astrocytes During Brain Development and Senescence. In: Omega Fatty Acids in Brain and Neurological Health. Elsevier, pp 479–490

79. Xiong Y, Li X, Liu J, Luo P, Zhang H, Zhou H, Ling X, Zhang M, Liang Y, Chen Q, Xing C, Li F, Miao J, Shen W, Zhou S, Wang X, Hou FF, Liu Y, Ma K, Zhao AZ, Zhou L (2024) Omega-3 PUFAs slow organ aging through promoting energy metabolism. Pharmacol Res 208:107384. 10.1016/j.phrs.2024.107384

